# Lifestyle and Genetic Factors Modify Parent-of-Origin Effects on the Human Methylome

**DOI:** 10.1101/2021.06.28.450122

**Authors:** Yanni Zeng, Carmen Amador, Chenhao Gao, Rosie M. Walker, Stewart W. Morris, Archie Campbell, Azra Frkatović, Rebecca A Madden, Mark J. Adams, Shuai He, Andrew D. Bretherick, Caroline Hayward, David J. Porteous, James F. Wilson, Kathryn L. Evans, Andrew M. McIntosh, Pau Navarro, Chris S. Haley

## Abstract

**Background:** parent-of-origin effects (POE) play important roles in development and complex disease and thus understanding their regulation and associated molecular and phenotypic variation are warranted. Previous studies have mainly focused on the detection of genomic regions or phenotypes regulated by POE. Understanding whether POE may be modified by environmental or genetic exposures is important for understanding of the source of POE-associated variation, but only a few case studies addressing these modifiable POE exist.

**Methods:** in order to understand this high order of POE regulation, we screened 101 genetic and environmental factors such as “predicted mRNA expression levels” of DNA methylation/imprinting machinery genes and early/late lifestyle/environmental exposures. POE-mQTL-modifier interaction models were proposed to test the potential of these factors to modify POE at DNA methylation using data from Generation Scotland: The Scottish Family Health Study(N=2315).

**Results:** a set of vulnerable/modifiable POE-CpGs were identified (modifiable-POE-regulated CpGs, N=3). Four factors, “lifetime smoking status” and “predicted mRNA expression levels” of *TET2*, *SIRT1* and *KDM1A*, were found to significantly modify the POE on the three CpGs in both discovery and replication datasets. Importantly, the POE on one of the CpGs were modified by both genetic and environmental factors. We further identified plasma protein and health-related phenotypes associated with the methylation level of one of the identified CpGs.

**Conclusions:** the modifiable POE identified here revealed an important yet indirect path through which genetic background and environmental exposures introduce their effect on DNA methylation, motivating future comprehensive evaluation of the role of these modifiers in complex diseases.

**Key Messages:**

- Previous population studies showed that parent-origin-effects(POE) on human methylome can be widespread and affect health-related traits and diseases.
- Whether the POE remained stable throughout the life or can be modified by genetic or environmental factors were largely unknown.
- By systematically screening 101 genetic and environmental factors in a large cohort(GS:SFHS) we provided the first population-level replicated evidence that those measuring lifestyle (smoking) and predicted expression of DNA methylation- or imprinting-machinery genes are amongst the factors that can modulate the POE of mQTLs for a set of CpG sites.
- We found those modifiable-POE-regulated CpGs are also phenotypically relevant – one is associated with the plasma levels of CLEC4C and health-related phenotypes such as HDL levels.
- The modifiable POE identified here revealed an important yet indirect path through which genetic background and environmental exposures introduce their effect on DNA methylation and their potential phenotypic consequences. This also provided a paradigm for further studies to explore how environmental and genetic effects can be integrated at methylation level.

## Introduction

Illustrating the sources of variation in DNA methylation lays the foundation for understanding epigenetic-based biomarkers for disease risk and progress prediction [1, 2]. DNA methylation is known to be influenced by additive and non-additive genetic and environmental factors [3–5]. As a special form of non-additive genetic effects, parent-of-origin effects (POE) on the human methylome manifest as differences in methylation levels between the reciprocal heterozygotes of the mQTL depending on the allelic parent-of-origin (Figure 1)[6]. Through selectively silencing the maternal or paternal allele, genomic imprinting has been considered as the major driving force creating the POE phenomenon [6]. We and others have shown that POE-influenced methylation sites are not rare, many are regulated by mQTLs, and that they follow one of the three classical imprinting patterns: parental, bipolar dominance and polar dominance (Figure 1) [4, 7, 8]. Although they only comprise a small proportion of the genome, POE (imprinting)-regulated CpGs and genes have been found to be important for developmental, metabolic and behavioral traits [4, 9].

**Figure 1.**
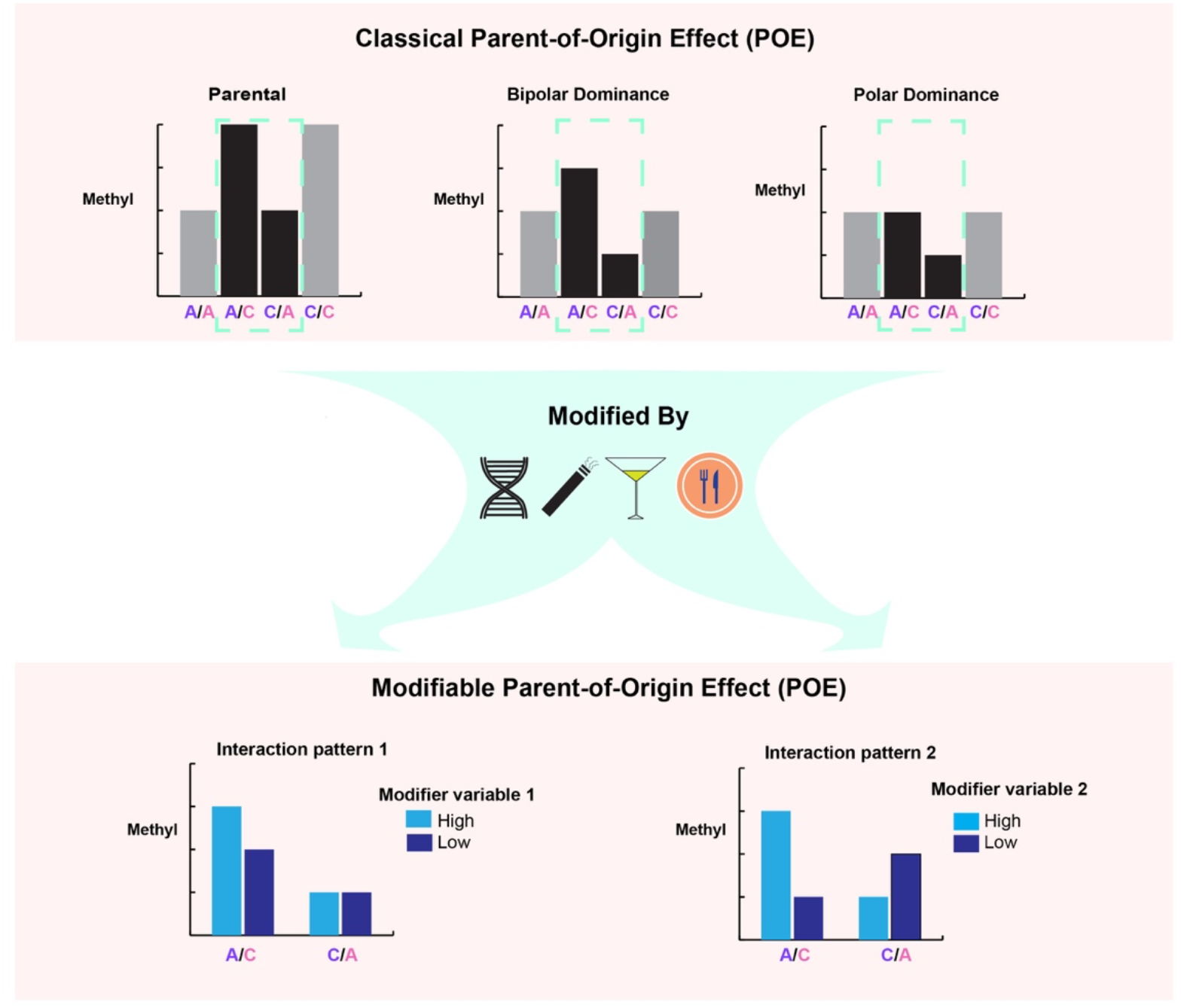
Patterns of classical and modifiable parent-of-origin effect (POE) regulation on DNA methylation. X axis: mQTL genotypes, left purple: paternal allele, right pink: maternal allele. Y axis: methylation level of the regulated CpG. Upper panel: classical POE patterns including parental and complex (dominance) POE patterns. Parental patterns show two levels of methylation depending on the expressed allele and the allelic effect. Complex POE manifests as the two homozygous group having the same methylation level whereas the heterozygous groups are different. Dashed box: difference between methylation level of heterozygous groups of the mQTL is the hallmark of POE. Bottom: scenarios when the POE is modified by genetic or environmental factors, leading to the alteration of POE for different levels of the modifier.

Despite their functional importance, the POE patterns in those POE (imprinting)-influenced regions can fluctuate as a consequence of genetic and environmental variation. Previous studies have reported that a large fraction of imprinted regions deviated from mono-allelic expression and that birth phenotypes were associated with the extent of this deviation [10]. A case study on an imprinting influenced long non-coding RNA, lncRNA nc886, found that the imprinting status of this locus is tunable by both genetic variants and environmental factors such as maternal nutrition and maternal age[11]. Importantly, the altered POE has phenotypic consequences: loss of imprinting of nc886 in infants at birth resulted in increased body mass index (BMI) in childhood [11]. For the majority of other POE-influenced regions, however, whether POE are modifiable under certain conditions remains unknown. Here we aim to explore modifiable POE, manifesting as the altered methylation difference between reciprocal heterozygotes of the mQTL due to effects from certain genetic or environmental modifiers (Figure 1), which potentially represents an important layer of POE-related regulation requiring systematic examination.

To search for modifiable POE on CpGs, a key question is which modifiers may have the potential to regulate the POE. Genomic imprinting, which likely underlies the POE, involves complex and multi-stage DNA methylation reprogramming processes, from the slow erasure of methylation at primordial germ cell stage, to the establishment of imprinted methylation signatures at germ cell stage, followed by pre-implantation maintenance of the imprinted methylation pattern during the global demethylation event, which is subsequently maintained post-implantation [12]. A number of gametic and zygotic genetic factors were found to be involved in these processes, such as those functioning in folate metabolism, the DNA methylation machinery (writers, erasers) and the proteins with which they interact [12]. Additionally, imprinting-related processes have also been found to be sensitive to environmental insults, such as the stress induced by assisted reproductive technologies, nutritional deficiency and adverse exposures [12–14]. Given that previous studies of modifiers of genomic imprinting were mostly case studies of individual factors, a systematic and population-wide scanning for genetic and environmental modifiers of POE is essential to fully characterize POE regulation.

In this study, we used Generation Scotland: The Scottish Family Health Study (GS:SFHS), a large family-based population cohort with extensive environmental and phenotypic records [15, 16], genome-wide genotypes and DNA methylation data [17, 18], to identify genetic and environmental factors that modify the POE on the human methylome (Figure 1). Figure 2 illustrates the study design. Based on the 2372 previously identified independent mQTL-CpGs pairs containing mQTLs with parent-of-origin effect (1895 independent mQTLs; 381211 SNPs in total) and their regulated CpGs (399 independent CpGs; 586 CpGs in total)[4], we proposed an interaction model which tests for significant interaction effects (Figure 1) between each of the 101 candidate environmental/genetic modifiers available in GS:SFHS and the parent-of-origin effect of each mQTL on the corresponding targeted CpG. Significant results from discovery samples (GS:SFHS set1, N=1663) were tested in replication samples (GS:SFHS set2, N=652). Plasma protein levels and health-related phenotypes associated with the modifiable-POE-regulated CpGs were also identified, suggesting phenotypic relevance for this special class of CpGs.

**Figure 2.**
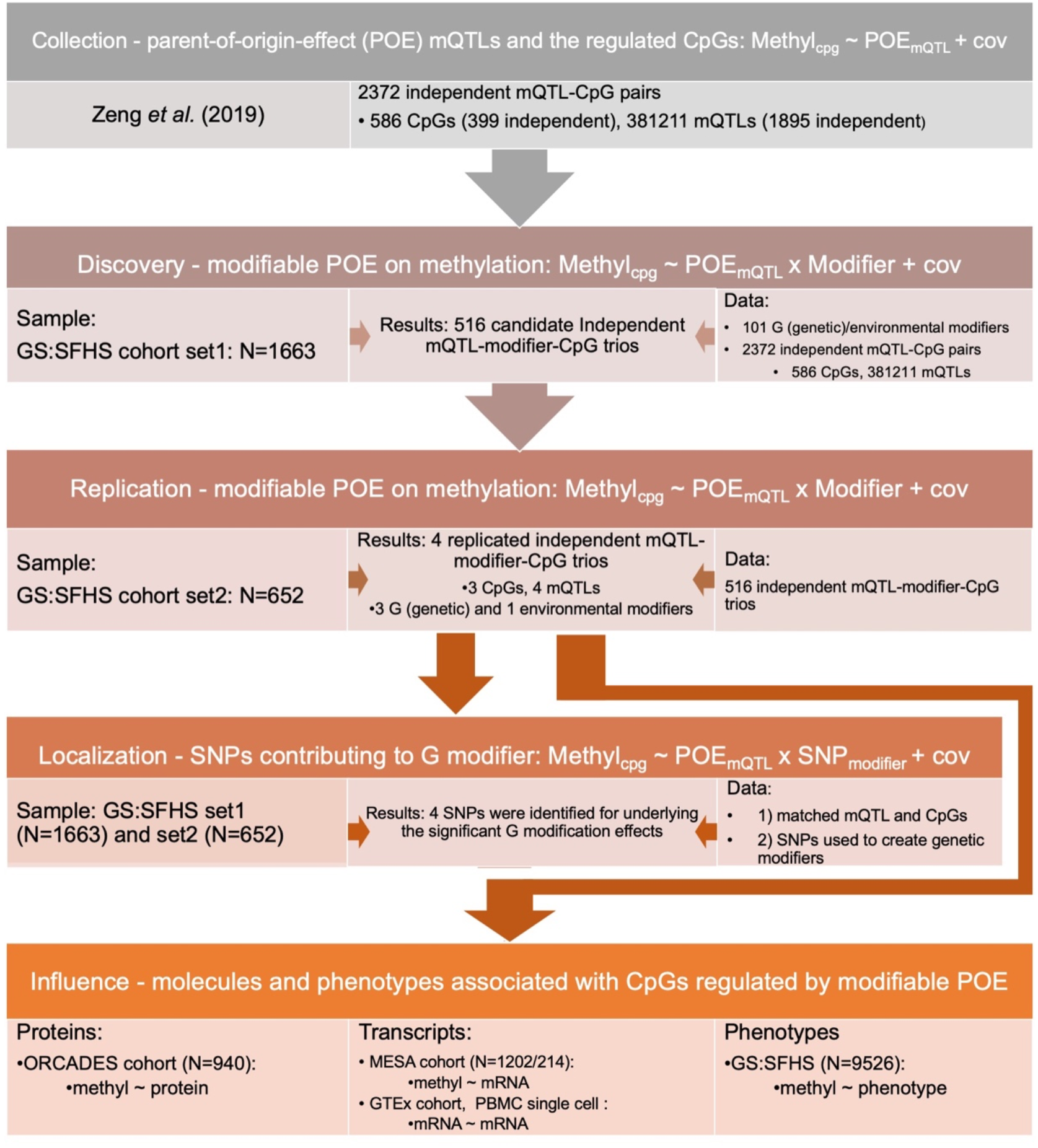
Design of the study. Cov: covariates fitted in the model. Zeng *et al.*(2019): the study which reported CpGs regulated by POE and the mQTLs that induce the POE for 586 CpGs (reference 4).

## Methods

### Population sample

Generation Scotland: the Scottish Family Health Study (GS:SFHS) is a deeply phenotyped population cohort [15, 16] with genome-wide genotypes available for 19994 participants, among which 9526 also have DNA methylation data available[15, 17, 18]. We used GS:SFHS to identify CpGs regulated by modifiable POE, and to explore phenotypes associated with those CpGs.

ORCADES is a family-based cross-sectional study which recruited 2078 participants between 2005 and 2011 from the Orkney Isles in northern Scotland [19]. Proteomic and DNA methylation data were available for a subset of 940 participants and were used here for association test between methylation sites and plasma protein levels (see below).

### GS:SFHS cohort: genotypes and inferences of parent-of-origin transmission of alleles in offspring

Genome-wide genotypes were generated using the Illumina Human OmniExpressExome -8- v1.0 array [20]. Phasing, imputation and quality control were described previously [4, 21]. In total, 7108491 high-quality imputed common SNPs (MAF ≥.01, info score ≥ 0.8) for 19994 participants were available for subsequent analyses. Among those individuals, there were 7139 offspring with at least one of their parents genotyped in GS:SFHS, which allowed us to successfully infer parent-of-origin allelic inheritance of all imputed common SNPs in 7106 offspring with high accuracy [4].

### GS:SFHS cohort: DNA methylation

In GS:SFHS, genome-wide DNA methylation data were produced through a related Stratifying Resilience and Depression Longitudinally (STRADL) project [18]. In 2016, the first wave of methylation data was generated on 5081 participants. These were used as discovery subset. 1663 of these participants also had imputed genotype information with parent-of-origin alleles successfully inferred and were used for the scanning for modifiable POE here. In 2019, another wave of methylation data was generated on an independent subset of 4445 participants. These data were used as replication subset. 652 out of these 4445 participants had imputed genotype information with parent-of-origin alleles successfully inferred. Based on a pipeline proposed previously [4], the two datasets were generated, processed and quality controlled in consistent way [22], which was briefly described in text s1.

### GS:SFHS cohort: Environmental/genetic modification variables

#### Environmental modifier variables

The core GS:SFHS cohort has rich collections of environmental variables[15]. Moreover, 98% of GS:SFHS participants gave informed consent for data linkage with historic Scottish birth cohorts which contain collections of birth and maternity information(Text s2). In total, we were able to collect 75 environmental variables and used them in downstream analyses. A full list of environmental variables is given in Table s1.

#### Genetic modifier variables

We considered two major sources of genetic modifiers for POE:

1. Predicted mRNA expression levels of 17 DNA methylation or imprinting-specific machinery genes imputed by PrediXcan [23].
2. Nine genetic risk scores for folate metabolism.

Details of those genetic modifiers were described in Text s2 and table s1,2.

### POE-mQTL-modifier interaction models

We applied a POE-mQTL-interaction model to test whether environmental or genetic factors could modify the POE induced by mQTL on CpGs. The model built on a linear regression model that we used to identify POE-specific mQTL-CpG pairs (mQTL with a parent-of-origin effect and the CpG it regulated) in our previous study [4, 24]:

#### model 1 - non-interaction model

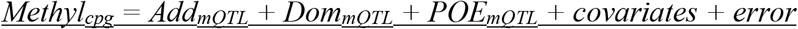

Where for the additive genetic variable (Add_mQTL_), the dominance genetic variable (Dom_mQTL_) and the parent-of-origin effect (POE_mQTL_), genotypes were coded as below:

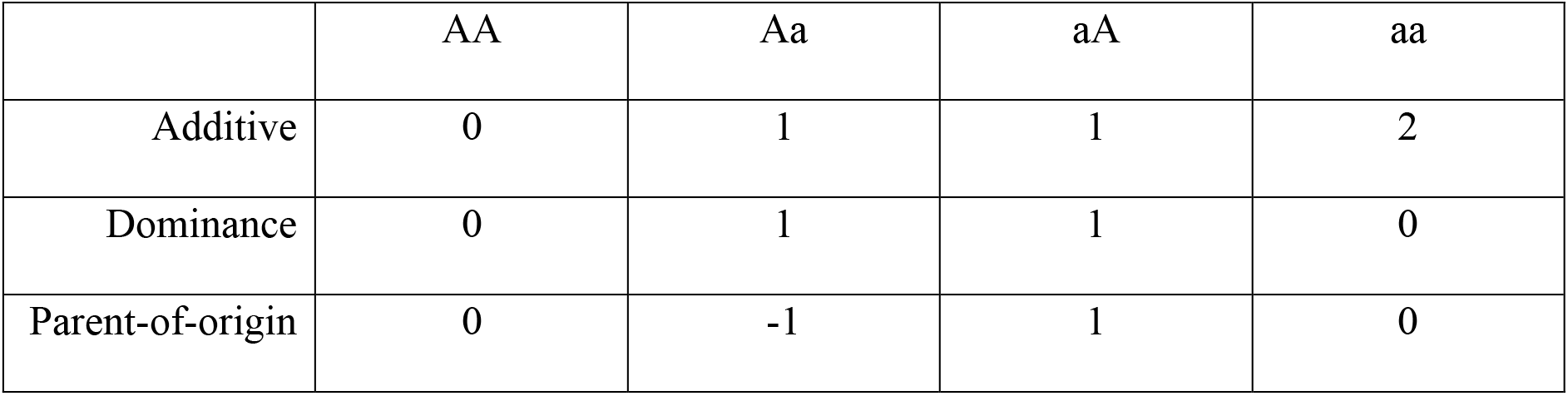

In this study, we applied an interaction model that additionally includes a modifier variable, *Mod*, and its interaction with the additive genetic effect and the POE:

#### model2 - interaction model

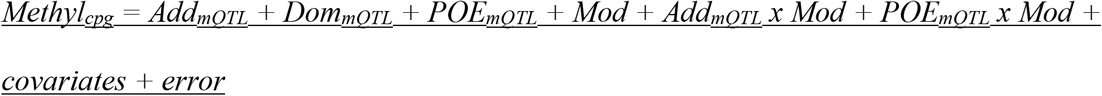

Where the modifier variable (*Mod*) was one of the environmental/genetic variables collected/derived as described in the section “GS:SFHS cohort: Environmental/genetic modification variables”. Covariates included age, sex, cell proportions, smoking variables (“pack years” and “lifetime smoking status”, which were not included as covariates when they were the tested modifier factor) and principal components (PCs) derived from an OMIC-relationship-matrix (ORM) created in OSCA [25] using all measured DNA methylation sites. To avoid removing genetic signals of interest by fitting ORM-PCs, we only included ORM-PCs among the top 20 ORM-PCs that were not significantly associated with any common SNP(determined through performing GWAS for ORM-PCs). We applied this model and tested the significance of the interaction effect between each mQTL’s POE and the modifier variable (*POE_mQTL_ x Mod*) on the corresponding CpG. The mQTL-CpG pairs tested were the 2372 independent POE-specific mQTL-CpG pairs which we reported previously [4]. The interaction between the additive effect and the modifier (*Add x Mod*) was jointly fitted in the model. For simplicity we did not fit an interaction between the dominance effect and the modifier (*Dom x Mod*) in the global scan, but we did this in sensitivity tests for the significant trios we identified. The results indicated only a minor contribution of the *Dom_mQTL_ x Mod* effect (Table s3). Multiple testing correction was performed by a combination of a global permutation test and mQTL-modifier-CpG trio-specific permutation tests for *POE_mQTL_ x Modifier* interaction effect at discovery stage(FDR≤0.05, details see Text s3, Table s4). A successful replication required to both reach statistical significance (FDR≤0.05) and patten consistency (details in Text s3, Table s4-6). Visualization of the results was performed using the R package coMET and ggplot2 [26, 27].

### Identification of proteins and mRNAs associated with CpGs regulated by modifiable POE

#### Association between DNA methylation and protein levels

##### ORCADES cohort-DNA methylation and proteomic data

DNA methylation was measured from whole blood samples using Illumina MethylationEPIC Array for 794627 CpG sites in 1052 samples (quality control and pre-correction in Text s4). Abundance of plasma proteins was measured from the fasted EDTA plasma samples for a subset of 1051 participants using Olink Proseek Multiplex cardiometabolic, cell regulation, cardiovascular 2 and 3, developmental, immune response, inflammation 1, metabolism, neuro exploratory, neurology, oncology and organ damage panels. The raw NPX values were used in analysis.

##### Association test between DNA methylation and plasma protein levels

A linear mixed model was used to compute methylation residuals after accounting for genetic structure in ORCADES by fitting a random effect represented in the genomic relationship matrix. To assess whether the CpGs (N_cpg_=3) significantly regulated by interaction effects (*POE_mQTL_ x Mod*) were associated with the abundance of any plasma protein (N_protein_=1092), the association between adjusted methylation of CpGs of interest with each measured protein was tested using a linear regression model in 940 participants where DNA methylation and proteomic data were simultaneously available:

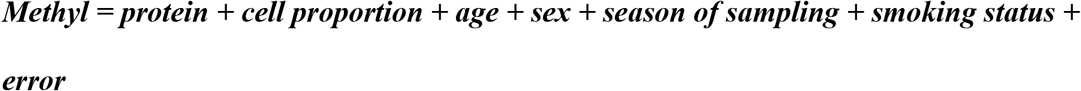

The Bonferroni method was applied to correct for multiple testing (N_correction_=3×1092=3276).

#### Correlations between DNA methylation vs mRNA levels

Transcriptomic and DNA methylation data from human peripheral monocytes and T cells in the Multi-Ethnic Study of Atherosclerosis (MESA) study were downloaded from the NCBI GEO database (Series GSE56047 and GSE56580) [28]. mRNA was measured using the Illumina HumanHT-12 v4 Expression BeadChip, DNA methylation levels were measured using the Illumina HumanMethylation450 BeadChip [28]. Quantile-normalized signal for mRNA (log2 transformed) and DNA methylation data (M-values) were simultaneously available for peripheral monocytes (CD14+) in 1202 participants and for peripheral T cells (CD4+) in 214 participants and were used to calculate the Spearman correlation between DNA methylation and mRNA.

#### Correlations between mRNAs levels

At the population-level, correlations between mRNA levels of target genes were calculated using the Spearman method on GTEx whole-blood data using the GEPIA portal [29]. At single cell level, a normalized single-cell matrix for 63628 peripheral blood mononuclear cell (PBMC) cells from a healthy donor were obtained from the website http://tisch.comp-genomics.org/gallery/. Feature counts for each cell were normalized by “LogNormalize”, a global-scaling method that normalizes the cellular feature expression by dividing the total counts for that cell, multiplied that by a scale factor (10000 by default), followed by a natural-log transformation. Spearman’s correlation between mRNA levels of target genes was calculated using cells where normalized expression levels of both genes were larger than zero.

### Phenome-wide association test for DNA methylation sites regulated by modifiable POE

We collected 79 phenotypes measured in GS:SFHS (recorded dataset and linked data) to identify phenotypes correlated with DNA methylation levels at CpG sites targeted by modifiable POE. The full list of phenotypes tested can be found in Table s7. The correlation was tested by regressing the adjusted M-values of methylation sites on each phenotype variable, covariates included age, age^2^, cell proportion, sex, top 20 ORM-PCs and smoking variables (“pack years” and “lifetime smoking status”. These were not included as covariates when they were the tested phenotype). The test was performed in the discovery and replication samples separately and meta-analyzed using a random effect model using the R package metafor [30]. The sample size varied depending on the number of missing samples for each specific phenotype (Table s8). The Bonferroni method was used for multiple testing correction (N_correction_=79*3=237).

## Results

### DNA methylation sites targeted by modifiable POE

In GS:SFHS, we utilized information on 75 environmental and 26 genetic variables to test if any of them significantly modified the parent-of-origin effects of the mQTLs on methylation level of the targeted CpGs, altering the methylation difference between reciprocal heterozygotes of the mQTLs (Figure 1). The environmental modifiers reflected the environments/events the participants have been exposed to or have experienced, including those measuring baseline non-genetic effects (sex, age), medication, lifestyle, socioeconomic status, birth-related phenotypes (measured before or after participants’ birth) and menarche/menopause-related events; the genetic modifiers were genetic factors known to be involved in DNA methylation and imprinting processes, including “predicted whole blood mRNA expression levels” of DNA methylation- or imprinting-specific machinery genes, and polygenic risk scores (PRSs) for folate-related metabolism (details in methods and Table s1).

In the discovery stage, a global permutation test was combined with mQTL-modifier-CpG trio-specific permutation tests to determine mQTL-modifier-CpG trios for significant *POE_mQTL_ x Modifier* interaction effect on DNA methylation(methods). In the replication stage, a successful replication required both statistical significance and pattern consistency(direction-of-effect) for both the main POE and the *POE_mQTL_ x Modifier* interaction effect in tested trios. In total, four mQTL-modifier-CpG trios reached significance in the discovery sample (*P_global_permutation(FDR_adjusted)_ < 0.05*, *P_trio_permutation(Bonferroni_adjusted)_ < 0.05*) and were successfully validated in the replication sample (Table 1, text s2, tables s4-6). These four trios involved one environmental modifier: “lifetime smoking status”; and three genetic modifiers: “predicted mRNA expression levels” of *SIRT1* (a gene that protects methylation at imprinted loci by directly regulating acetylation of DNMT3L), *TET2* (a DNA demethylation gene) and *KDM1A* (a gene involved in removal of methylation and histone H3K4 in imprinted genes). Three CpGs, cg18035618, cg21252175, and cg22592140 were the methylation sites affected by these modification effects (Table 1), with the *POE_mQTL_ x Modifier* interaction effect explaining between 1.0% to 2.2% of their methylation variation (Figure 3a, Table s3). These CpGs displayed intermediate methylation levels, when compared with the less-peaked distribution of methylation levels of POE-regulated CpGs not influenced by modifiable effects, or the bi-directionally distributed methylation level of genomic CpGs not influenced by POE (Figure 3b). One of the three CpGs, cg18035618, was simultaneously targeted by both environmental and genetic modification effects (Table 1).

**Table 1.** Genetic/environmental modifiers displaying a significant interaction with the POE(mQTL) and the CpGs the interaction effects affect. CpG-POS/mQTL-POS: genomic position of the CpG and mQTL(hg19). Env: environmental modifier. dis: discovery sample, rep: replication sample. N: sample size. P(dis)/P(rep) and Est(dis)/Est(rep): P values and coefficients for the *POE_mQTL_ x Modifier* effect in the interaction model in discovery/replication samples. P(trio-specific-perm): P value for the *POE_mQTL_ x Modifier* effect from the trio-specific permutation test. Genomic position of the significant genetic modifiers: *KDM1A*: chr1: 23345941-23410182; *TET2*: chr4: 106067,032-106200973; *SIRT1*: chr10: 69644427-69678147.

| mQTL-CpG-modifier trio information |  |  |  |  |  |  |  | Discovery |  |  |  |  | Replication |  |  |  |
| --- | --- | --- | --- | --- | --- | --- | --- | --- | --- | --- | --- | --- | --- | --- | --- | --- |
| CpG: id | CpG-chr | CpG-POS | mQTL:<br>id | mQTL-chr | mQTL-POS | Modifier | Modifier<br>type | P(dis) | Est(dis) | N(dis) | P<br>(trio-specific<br>-perm) | Adjusted P<br>(trio-specific-perm) | P(rep) | Est(rep) | N(rep) | Adjusted<br>P(rep) |
| **cg22592140** | 7 | 130132419 | rs142053217 | 7 | 130258926 | mRNA expression of KDM1A | Genetic | 1.35E-06 | 6.35 | 1607 | 1.00E-05 | 1.09E-02 | 6.09E-04 | 6.52 | 645 | 2.79E-02 |
| **cg21252175** | 15 | 64973903 | rs111834387 | 15 | 64062211 | mRNA expression of TET2 | Genetic | 1.54E-06 | 3.77 | 1607 | <1e-5 | < 0.01 | 7.76E-11 | 13.80 | 647 | 2.43E-08 |
| **cg18035618** | 20 | 57415978 | rs117020124 | 20 | 57347435 | Lifetime smoking status | Env | 9.94E-07 | -0.04 | 1607 | <1e-5 | < 0.01 | 1.42E-03 | -0.05 | 647 | 4.85E-02 |
|  |  |  | rs117823351 | 20 | 57369773 | mRNA expression of SIRT1 | Genetic | 3.68E-10 | -0.24 | 1607 | <1e-5 | < 0.01 | 1.01E-03 | -0.22 | 646 | 3.95E-02 |

**Figure 3.**
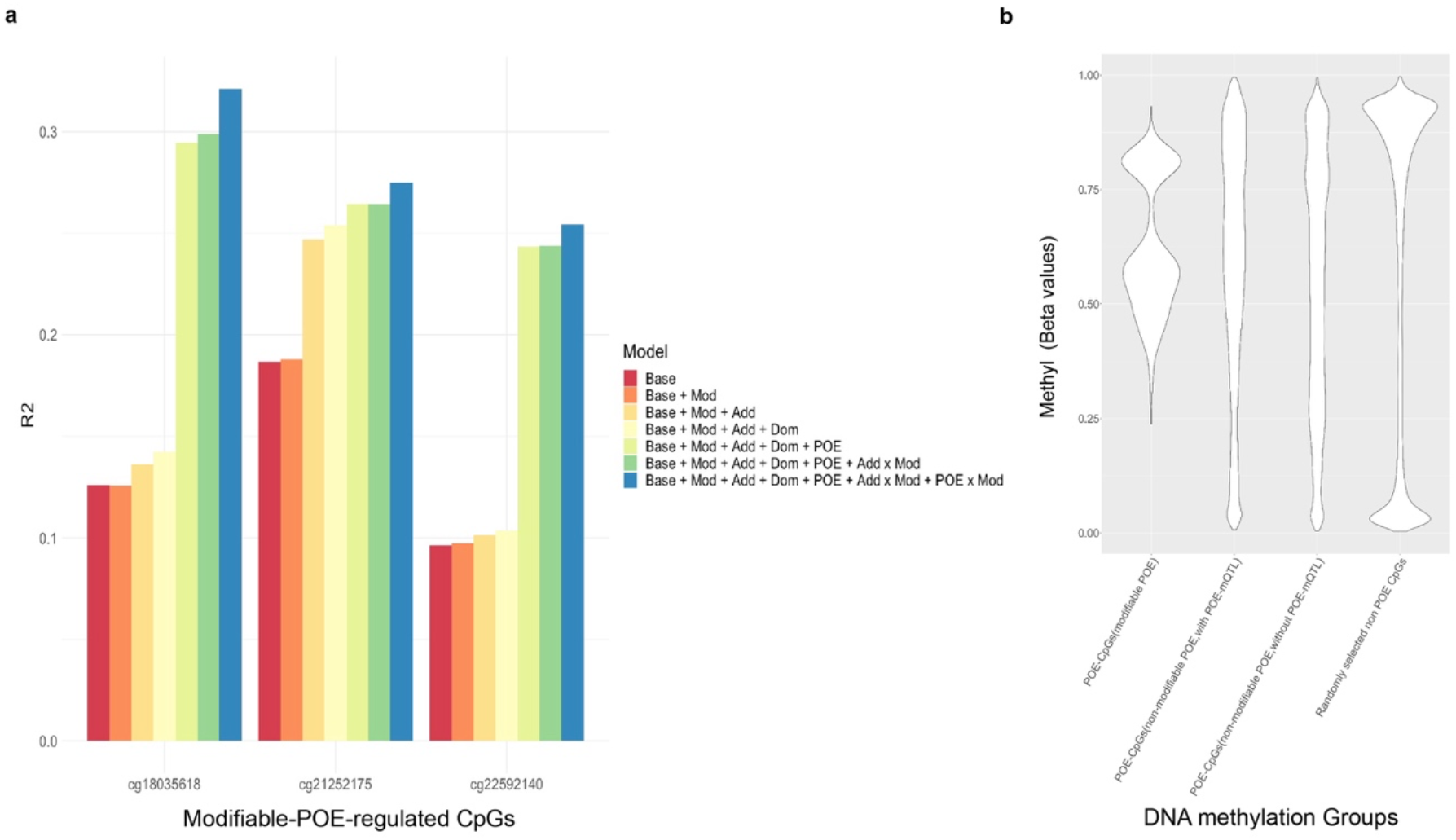
CpGs regulated by the modifiable-POE. a. Proportion of methylation variation explained by different models for the three CpGs regulated by modifiable-POE. The “Base model” accounted for age, sex, cell proportion, smoking variables (“pack years” and “lifetime smoking status”. These were not included as covariates when they were the tested modifier factor) and principal components of the OMIC (DNA methylation)-relationship-matrix. Mod: modifier; Add: additive genetic effect, Dom: dominance genetic effect. Add x Mod: interaction between additive genetic effect and the modifier; POE x Mod: interaction between parent-of-origin genetic effect and the modifier. b. Distributions of methylation levels of CpGs regulated by modifiable POE (N_CpG_=3), CpGs regulated by POE from known mQTLs but the POE is not modifiable (N_CpG_=583), CpGs regulated by POE but without an mQTL identified (the CpGs were reported in ref 11: Zeng *et al.* (2019), N_CpG_=398) and randomly selected non-POE CpGs from all QCed array probes (N_CpG_=10,000). Unrelated individuals from the GS:SFHS discovery subset were used to produce the plot.

cg18035618 (hg19: chromosome 20: 57415978) is located in an intron of the gene *GNAS* (Figure 4). The methylation level of cg18035618 was significantly modulated by a *POE_mQTL_ x Modifier* interaction between its mQTL, rs117020124, and “lifetime smoking status” (*P_dis_=*9.94×10^−7^, *P_rep_=*1.42×10^−3^, figure 4a, table 1), with current smokers displaying larger contrast in methylation levels of cg18035618 between heterozygous groups of the mQTL when compared with ex-smokers and never-smokers (Figure 5a). For this CpG a significant *POE_mQTL_ x Modifier* interaction was also detected between another independent mQTL, rs117823351, and “predicted mRNA expression levels of *SIRT1*” (*P_dis_*=3.68×10^−10^, *P_rep_=*1.01×10^−3^, figure 4b, table 1. *SIRT1* is located in chromosome 10), with lower *SIRT1* expression corresponding to an increased contrast of methylation levels between heterozygous groups of the mQTL (Figure 5b).

**Figure 4.**
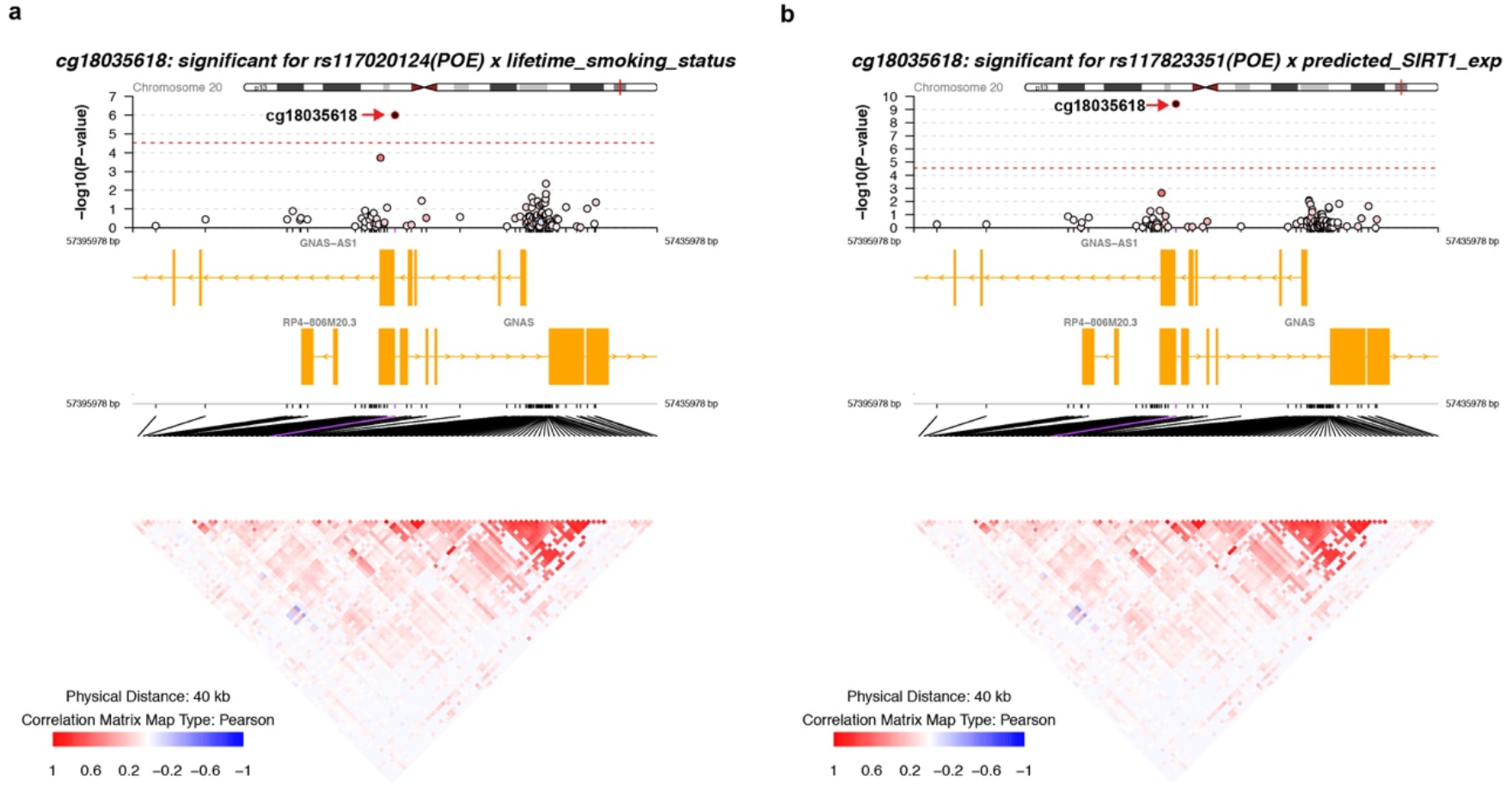
Regional plot of the modifiable-POE affecting cg18035618 and nearby CpGs within a 20kb distance. a. Interaction effect between the POE of cg18035618’s mQTL rs117020124 and “lifetime smoking status”; b. Interaction effect between the POE of cg18035618’s mQTL rs117823351 and ‘predicted mRNA expression levels of *SIRT1’*. Upper panel: −log10 (P-value): minus log10 P-value of the *POE_mQTL_ x Modifier* interaction effect. Dots show nearby measured CpGs located within a 20kb distance from cg18035618, filling colour represents the correlation of methylation levels with cg18035618: red: positive correlation; blue:negative correlation; white: no significant correlation.. Middle panel: genes located within the 40kb genomic region. Bottom panel: correlation matrix between CpGs.

**Figure 5.**
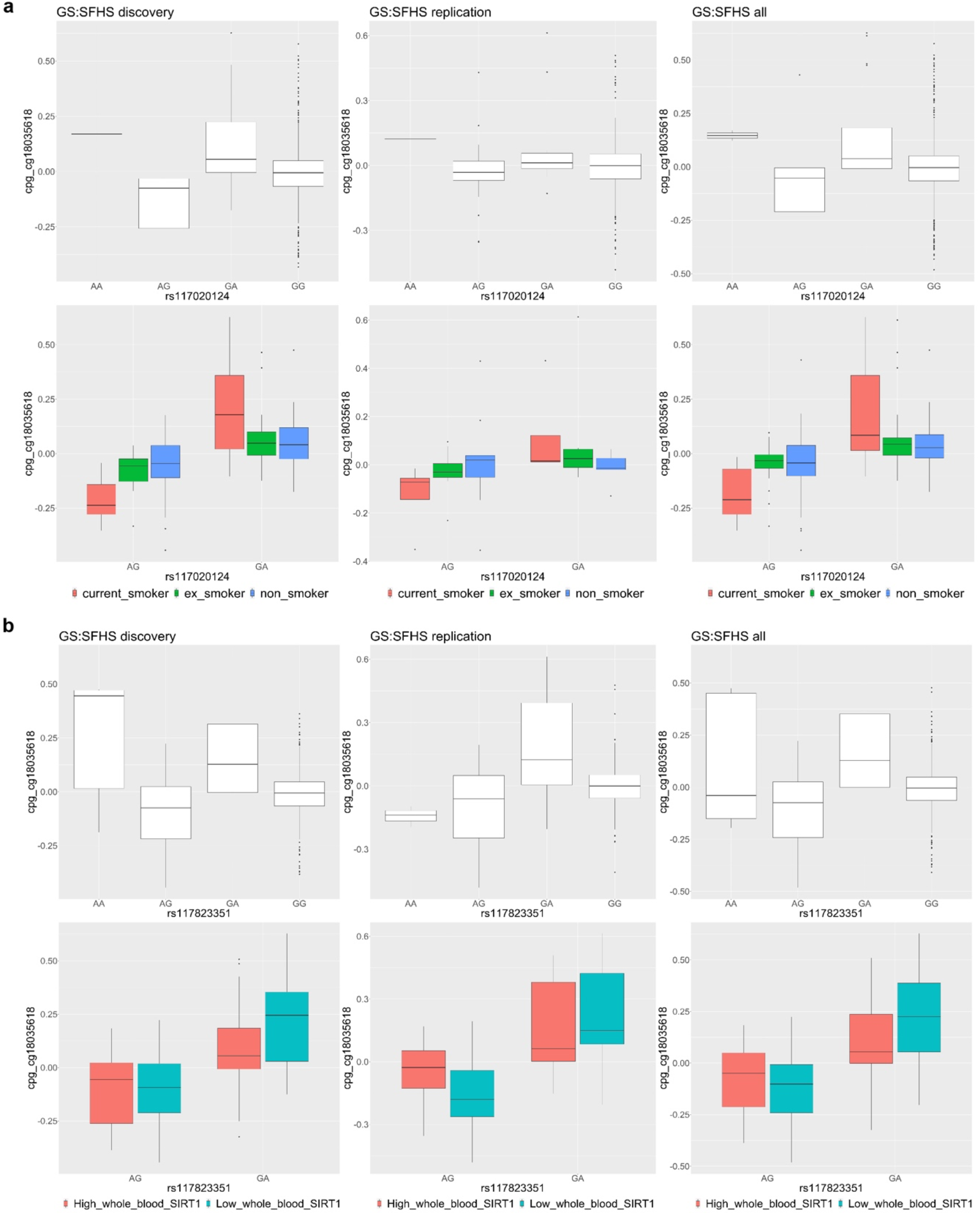
Both environmental and genetic factors significantly modified the POE of mQTLs on cg18036618. a.top:cg18036618 was regulated by the POE of the mQTL rs117020124. bottom: the POE of rs117020124 was modified by lifetime smoking status. The contrast in methylation levels of cg18035618 between heterozygotes of the mQTL rs117020124 is largest in current smoker group. b.top:cg18036618 was regulated by the POE of the mQTL rs117823351. bottom: the POE of rs117823351 was modified by “predicted mRNA expression level of *SIRT1*”. The contrast in methylation levels of cg18035618 between heterozygotes of the mQTL rs117823351 is larger in individuals with low SIRT1 expression.

To rule out the possibility that the sharing of the methylation target (cg18035618) by both genetic and environmental modifiers was due to genetically influenced environmental effects (i.e., *SIRT1* expression influences smoking status) [31], we calculated the correlation between “lifetime smoking status” and “predicted mRNA expression levels of *SIRT1*” and found no significant correlation (R=0, *P*=0.968).

cg21252175 (hg19: chromosome15: 64973903) is located in the 3’ UTR region of gene *ZNF609* (Figure s1a). For this CpG, a significant *POE_mQTL_ x Modifier* interaction was detected between the mQTL rs111834387 and “predicted mRNA expression levels of *TET2*” (*P_dis_=*1.54×10^−6^, *P_rep_*=7.76×10^−11^. *TET2* is located in chromosome 4), with lower expression of *TET2* reducing the contrast in methylation levels of cg21252175 between heterozygous groups of the mQTL (Table 1, figure s2a). cg22592140, a CpG located in an intron of gene *MEST* (Figure s1b), was significantly regulated by a *POE_mQTL_ x Modifier* interaction between its mQTL rs142053217 and “predicted mRNA expression levels of *KDM1A*” (*P_dis_*=1.35×10^−6^, *P_rep_*=6.09×10^−4^. *KDM1A* is located in chromosome 1), with lower expression of *KDM1A* increasing the contrast in methylation levels of cg22592140 between heterozygous groups of the mQTL (Table 1, figure s2b).

### Localization of regulatory SNPs contributing to the genetic modification effect

Considering that the three identified genetic modifiers (“predicted mRNA expression levels” of *SIRT1, TET2, KDM1A*) are essentially weighted combinations of allelic scores at multiple regulatory SNPs, we next tested whether the genetically driven modification effects we detected here can be recapitulated by the *POE_mQTL_ x SNP* interactions between mQTLs and the SNPs used to drive the significant genetic modifiers. Since “predicted mRNA expression” of *SIRT1*, *KDM1A* and *TET2* was derived from two, one and two SNPs respectively by the MASHR method in PrediXcan(Methods)[32], we tested *POE_mQTL_ x SNP* interactions for those 5 SNPs (N_test_=5). We identified four of the five SNPs significantly interacting with the POE_mQTL_, that is, regulating the CpGs where the interaction effect was initially detected (Table 2, Figure 6). For example, for cg18035618 we detected a significant interaction effect (*P_dis_*=3.69×10^−9^, *P_rep_*=4.33×10^−3^) between rs932658, a SNP used in the prediction model for *SIRT1* expression, and rs117823351, the mQTL that significantly interacted with “predicted mRNA expression levels of *SIRT1*”. When accounting for these significant *POE_mQTL_ x SNP* interaction effects as conditional items in the interaction model for the three genetic-modifiers, the interaction effect at the genetic-modifier-level (*POE_mQTL_ x Modifier_Genetic_*) were reduced to a non-significant level, suggesting the leading role of *POE_mQTL_ x SNP* underlying the significant interaction effect from genetic modifiers we detected here (Table s10).

**Table 2.**
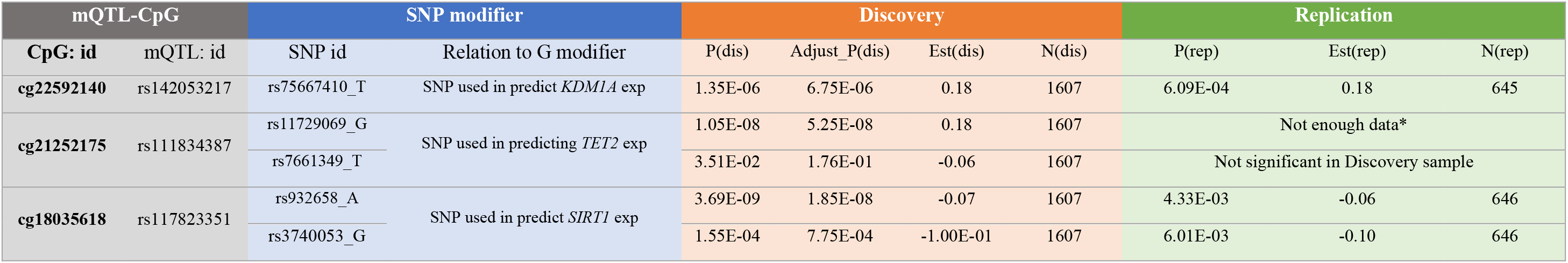
SNPs significantly interact with POE (*POE_mQTL_ x SNP*) and the regulated CpGs dis: discovery sample, rep: replication sample. N: sample size. P(dis)/P(rep) and Est(dis)/Est(rep): P values and coefficients for the POE_mQTL_ x SNP interaction in the interaction model in discovery/replication samples. *, due to the relatively small minor allele frequency (MAF) of rs111834387(MAF=0.01) and the limited sample size of replication sample, there was not enough data for this test in replication samples. Number of individuals within heterozygous groups available for testing *POE(rs111834387) x SNP(rs11729069)* interaction effect on cg21252175 in replication samples were shown in Table s9.

**Figure 6.**
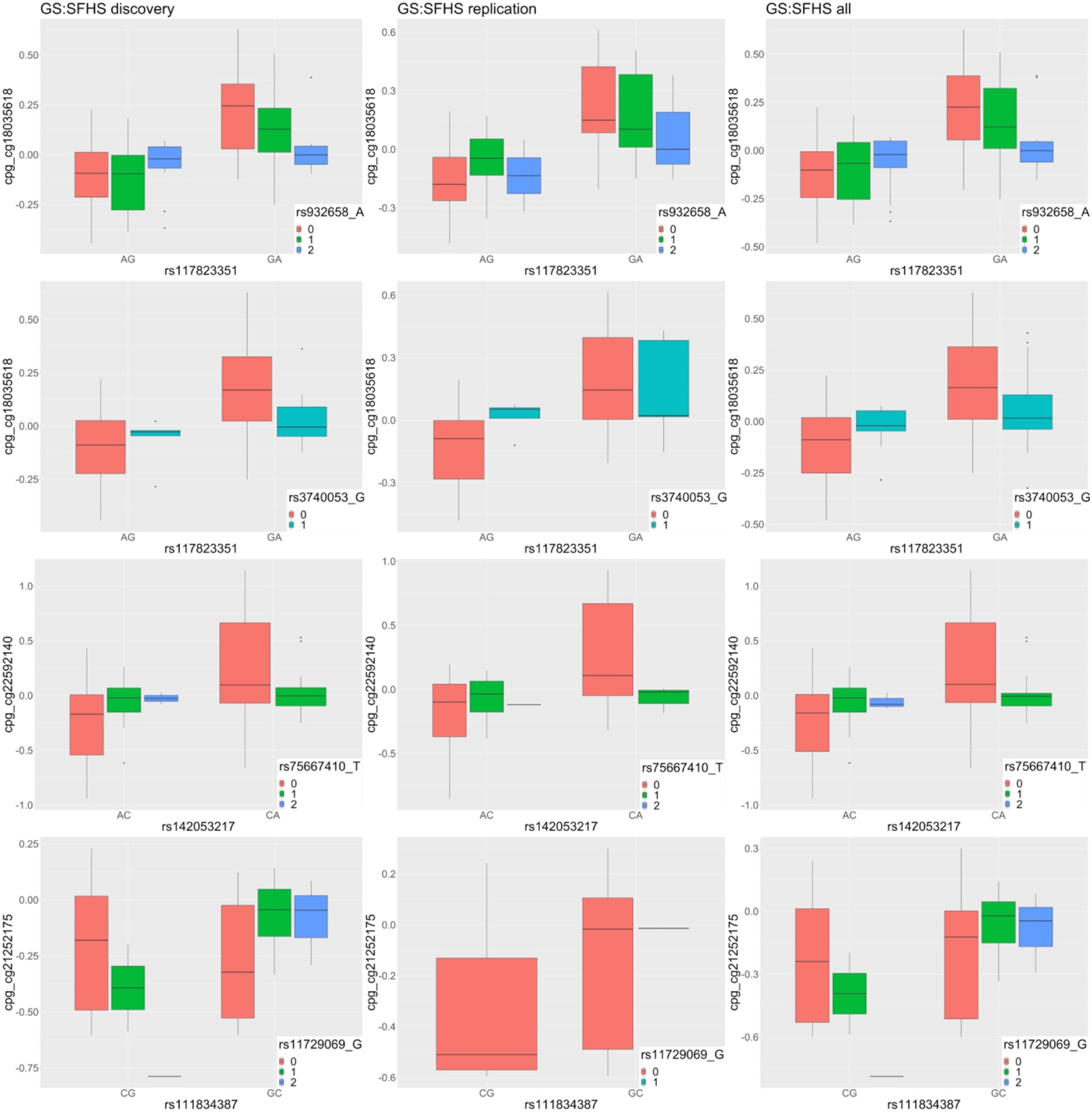
The three modifiable-POE-targeted CpGs were also significantly regulated by *POE_mQTL_ x SNP* interaction effects between the mQTLs and the SNPs used to drive the genetic modifiers. The contrast in methylation levels of the candidate CpGs in mQTL heterozygotes varied depending on the allelic dosage of the SNP used to derive the corresponding genetic modifier. *due to the limitation of sample size, minor homozygous/heterozygous genotype groups were missing in some tests.

### Proteins associated with modifiable-POE-regulated methylation sites

Association tests were performed using DNA methylation and proteomic data from ORCADES cohort (N_sample_=940, N_total_protein_=1092) to identify proteins associated with the three CpGs regulated by the modifiable POE. Only one CpG-protein pair passed Bonferroni-based multiple testing correction (N_test_=3,276, full results in Table s11): cg21252175 was significantly associated (in *trans*) with plasma protein level of CLEC4C, a protein from the Olink immune-response panel (Beta=0.049, *P_adj_=0.002*).

Since cg21252175 is located in the UTR3 of gene *ZNF609*, we further examined whether the association between cg21252175 and CLEC4C protein levels implied a link between *ZNF609* and *CLEC4C*. Using data from MESA study [28], a significant and positive correlation was detected between methylation levels of cg2125217 and mRNA levels of *ZNF609* in both CD4+ peripheral T cells (R=0.15, *P*=0.03) and CD14+ peripheral monocytes (R=0.15, *P*=1.52×10^−6^, figure s3a). Using whole blood data from the GTEx consortium through the GEPIA portal[29, 33], mRNA expression level of *ZNF609* was significantly correlated with mRNA expression levels of *CLEC4C* at population level (R=0.21, *P*=1.1×10^−4^, figure s3b). Using a single-cell RNA-seq data of PBMC in an adult donor, mRNA expression levels of *ZNF609* and *CLEC4C* were significantly correlated at the cellular level (R=0.36, *P*=0.0002, figure s3c).

### Phenotypes associated with modifiable-POE-regulated methylation sites

To explore the association between variation in CpGs targeted by modifiable POE and heath/disease-related phenotypes, we collected 79 phenotypes in GS:SFHS (Table s7). Phenome-wide association tests relating methylation levels to phenotypes were performed for the three identified modifiable-POE-regulated CpGs using the whole GS:SFHS methylation sample (meta-analyzed across discovery (N_sample_=5081) and replication (N_sample_=4445) samples; N_sample_=9526.). After Bonferroni-based multiple testing correction (N_test_=79×3=237), two CpG-phenotype associations reached phenome-wide significance: cg21252175 was both associated with “lifetime smoking status” (*P_adj_*= 9.0×10^−5^) and high-density lipoprotein (HDL) levels (*P_adj_* =0.006) (Figure 7, Table s8). Although limited by sample size(Figure 7), cg21252175 was also associated with gestational age (measured as weeks at birth) at an adjusted P ≤ 0.06 level (*P_adj_*=0.056). These associations displayed consistent patterns across discovery and replication samples (Figure 7, Table s8).

**Figure 7.**
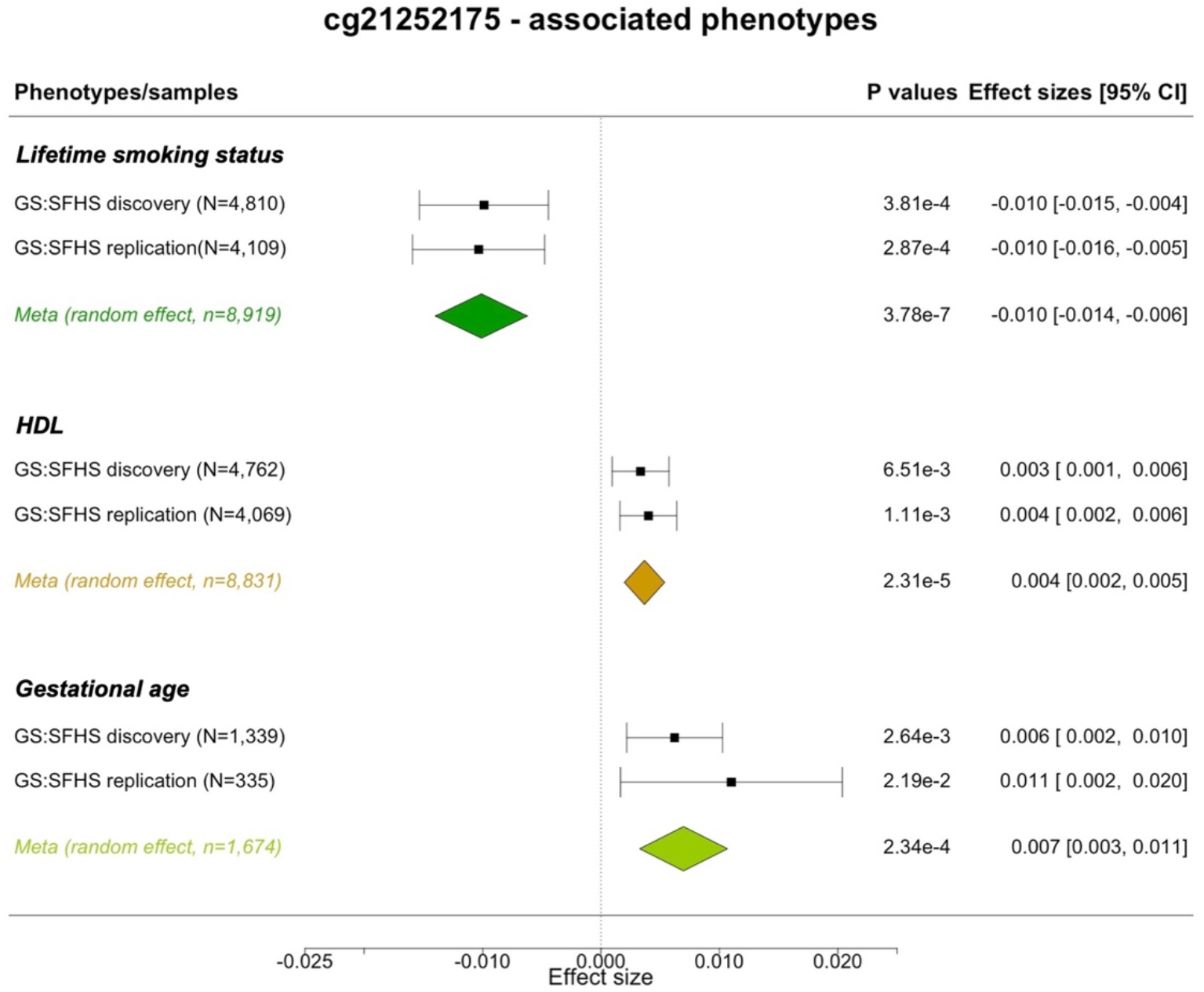
Forest plot for phenotypes associated with cg21252175. HDL: high-density lipoprotein. Meta: Meta-analysis performed using the random effect model.

## Discussion

In this study, we reported significant and replicated modification effects from both genetic and environmental variables on the parent-of-origin effect that affects DNA methylation levels at three CpGs. Identified environmental modifiers included “lifetime smoking status”; identified genetic modifiers included “predicted mRNA expression levels” of several DNA methylation/imprinting machinery genes (*SIRT1, TET2, KDM1A)*. Importantly, we found that both genetic and environmental modifiers were targeting the same CpG (cg18035618). These provided evidence for a special type of CpGs in the human genome regulated by the parent-of-origin-effects that are modulated by genetic or environmental modifiers. We further found that these CpGs are likely to be phenotypically relevant: at the molecular level, DNA methylation level at one the CpG, cg21252175, was associated with protein levels of the immune-response-related protein CLEC4C. At the phenotypic level, this CpG was associated with “lifetime smoking status”, HDL levels and gestational age(the latter at the suggestive significance level).

Statistically, the model we proposed here for detecting modifiable POE is built on a previous POE-mQTL model which we used to localize regulatory mQTLs for the POE-influenced CpGs (model 1 in Methods) [4, 24]. For those CpGs, one of the hallmarks of the POE-regulation was the methylation difference between the two heterozygous mQTL genotype groups (Figure 1)[4, 6]. Here, our new interaction model tests whether that difference remains stable or varies under certain conditions, that is, if the parent-of-origin effects are the same or different across different environments or in different genetic backgrounds. Biologically, this implies the existence of a new and different layer of regulation for DNA methylation: genetic/environmental factors could influence the level of DNA methylation of CpGs, not only through direct effects, but also through interacting with the POE (Figure 1).

Our results support this hypothesis. Smoking has been widely studied for its direct influence on DNA methylation [34, 35] and its interactions with additive genetic effects on methylation levels [36]. Here, for the first time, our study reported that smoking could also affect DNA methylation variation indirectly through interaction with a non-additive genetic effect, POE. Similarly, variation in DNA methylation and imprinting machinery genes, either in the forms of variable expression or mutation, have been known to directly affect DNA methylation. SIRT1 regulates DNA methylation and protects methylation at imprinted loci by directly regulating acetylation of DNMT3L [12, 37]. TET2 promotes DNA demethylation by converting 5-methyl-cytosine to 5-hydroxymethyl-cytosine and is required for demethylation at imprinted loci in the germline [12, 38]. KDM1A removes methylation of histone H3K4 in imprinted genes, its deficiency is associated with alterations in DNA methylation and expression at imprinted genes [12, 39]. Here, for the first time, we showed that besides known direct effects, these genetic factors also introduce indirect regulatory effects on DNA methylation levels through interactions with POE.

The detection of interactions between genetic modifiers and POE led us to further identify significant and replicated *POE_mQTL_ x SNP* interaction effects between mQTLs and SNPs used in imputing genetic modifiers. One important feature of the genetic modifier variables we derived here is that they represent the proportion of mRNA variation determined by germline genetic variation, which is constant and stable throughout the life [40]. Our results demonstrated that an individual’s genetic background of DNA methylation and imprinting machineries has the potential to modify POE. The localization of the genetic-based modification effect at regulatory SNPs of these DNA methylation and imprinting machinery genes strongly supports this, and importantly, indicated that genetic variation in machinery genes is an important source of epistasis. One of the machinery genes, *SIRT1*, has been well known for its role in mental disorders such as depression[41], but very few studies have examined its role as a modifier for non additive genetic effect such as POE. Our result revealed a new potential path of variation in this gene to introduce molecular differences.

The reason why POE can ever be modified is deeply rooted in the unique nature of genomic imprinting: established at an early developmental stage, needing to be protected from global-demethylation and maintained throughout the lifespan [12]. These complex and multi-stage processes have been shown to be vulnerable to environmental fluctuations and involve delicate regulation of multiple gametic and zygotic genetic factors, including *TET2*, *SIRT1*, *KDM1A* as we identified here.[11, 12]. Indeed, the vulnerability of at least a subset of POE-CpGs was revealed here, as the *POE_mQTL_ x Modifier* interaction effect explained a non-negligible proportion of methylation variance (1%-2.2%), and that at least one (cg18035618) of the three CpGs identified was targeted by independent environmental and genetic modifiers. Molecularly, we found one of the modifiable-POE influenced CpG (cg21252175) was associated with CLEC4C, an immune-response transmembrane protein treated as a marker gene for plasmacytoid dendritic cells [42]. Phenotypically, cg21252175 was associated with “lifetime smoking status”, HDL levels and gestational age (the latter at the suggestive significance level) (Figure 7) in our analysis, and was previously found to be associated with maternal body mass index/overweight/obesity [43]. These convergent lines indicated that this newly uncovered class of vulnerable POE-CpGs may play an important role in connecting early life stress, variations in genetic background and later life health issues.

There are limitations in this study. First, mRNA expression levels used here were predicted and only reflect genetically influenced expression variation (and not necessarily all of it). Future studies examining measured mRNA expression will be necessary to account for the modification effect from the environmentally determined fraction of mRNA expression. Second, since the analyses performed here simultaneously require DNA methylation data and SNP data with information of parent-of-origin of the alleles transmitted to the offspring, the sample size was relatively limited in this study. This is a proof of principle study and larger sample sizes will be needed to uncover more modifiable POE. Finally, the exact time window or developmental stage in which each environmental modifier exerted their influences remains unknown. Cohort data with higher resolution of environmental exposure records, in particular those measuring early life exposures, will be crucial to understand the vulnerable stage or stages for CpGs influenced by modifiable POE.

## Conclusions

we provided the first population-level evidence for modification effects from multiple genetic and environmental factors on parent-of-origin-effects at the DNA methylation level. A subset of parent-of-origin-effect-influenced CpGs that are vulnerable to modification effects were uncovered, which opens new questions for future profiling of the modification patterns and phenotypic consequences of this class of CpGs.

## Ethics approval

This study was ethically approved by the Tayside Research Ethics Committee (reference 05/S1401/89). Participants all gave written consent after having an opportunity to discuss the research and before any data or samples were collected.

## Data Availability Statement

Generation Scotland data are available from the MRC IGC Institutional Data Access / Ethics Committee for researchers who meet the criteria for access to confidential data. Generation Scotland data are available to researchers on application to the Generation Scotland Access Committee. The managed access process ensures that approval is granted only to research which comes under the terms of participant consent which does not allow making participant information publicly available.

## Acknowledgements

We want to acknowledge support from Genetics Core Laboratory at the Wellcome Trust Clinical Research Facility (Edinburgh, Scotland) for genotyping of the GS samples. We are grateful to all the families who took part, the general practitioners, and the Scottish School of Primary Care for their help in recruiting them, and the whole Generation Scotland team, which includes interviewers, computer and laboratory technicians, clerical workers, research scientists, volunteers, managers, receptionists, healthcare assistants and nurses. We would like to acknowledge the invaluable contributions of the research nurses in Orkney, the administrative team in Edinburgh and the people of Orkney. MESA and the MESA SHARe projects want to acknowledge the supported from the National Heart, Lung, and Blood Institute (NHLBI) in collaboration with MESA investigators. YZ wants to acknowledge support from Dr. Lucija Klaric for pre-processing the ORCADES data.

