## supplementary figures and texts for "Lifestyle and Genetic Factors Modify Parent-of-Origin Effects on the Human Methylome"

Figure s1. The regional plot for modifiable-POE-targeted CpGs cg21252175 and cg22592140.

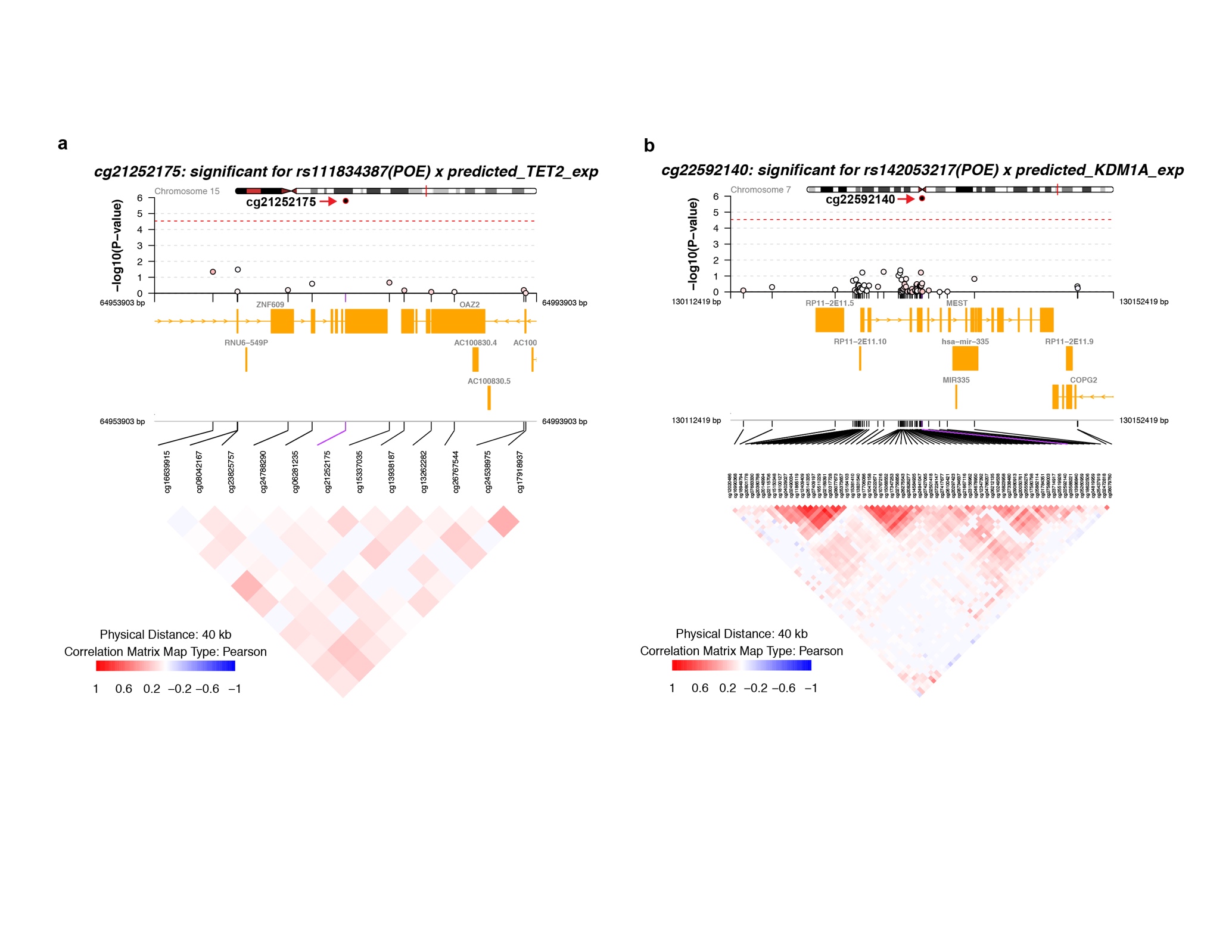

Figure s2 Genetic factors signficantly modified the POE of mQTL on cg21252175(a) and cg22592140(b).

a

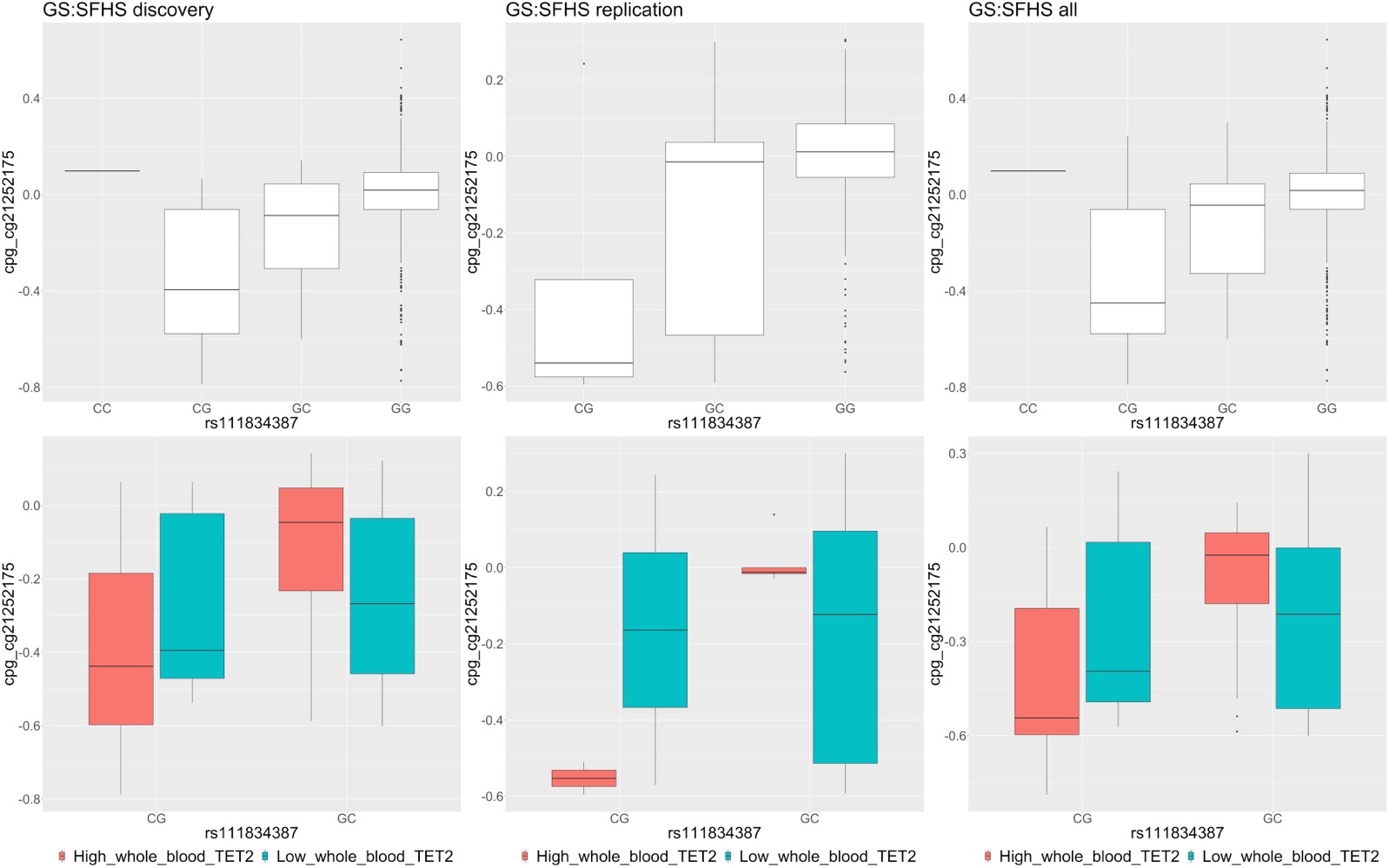

b

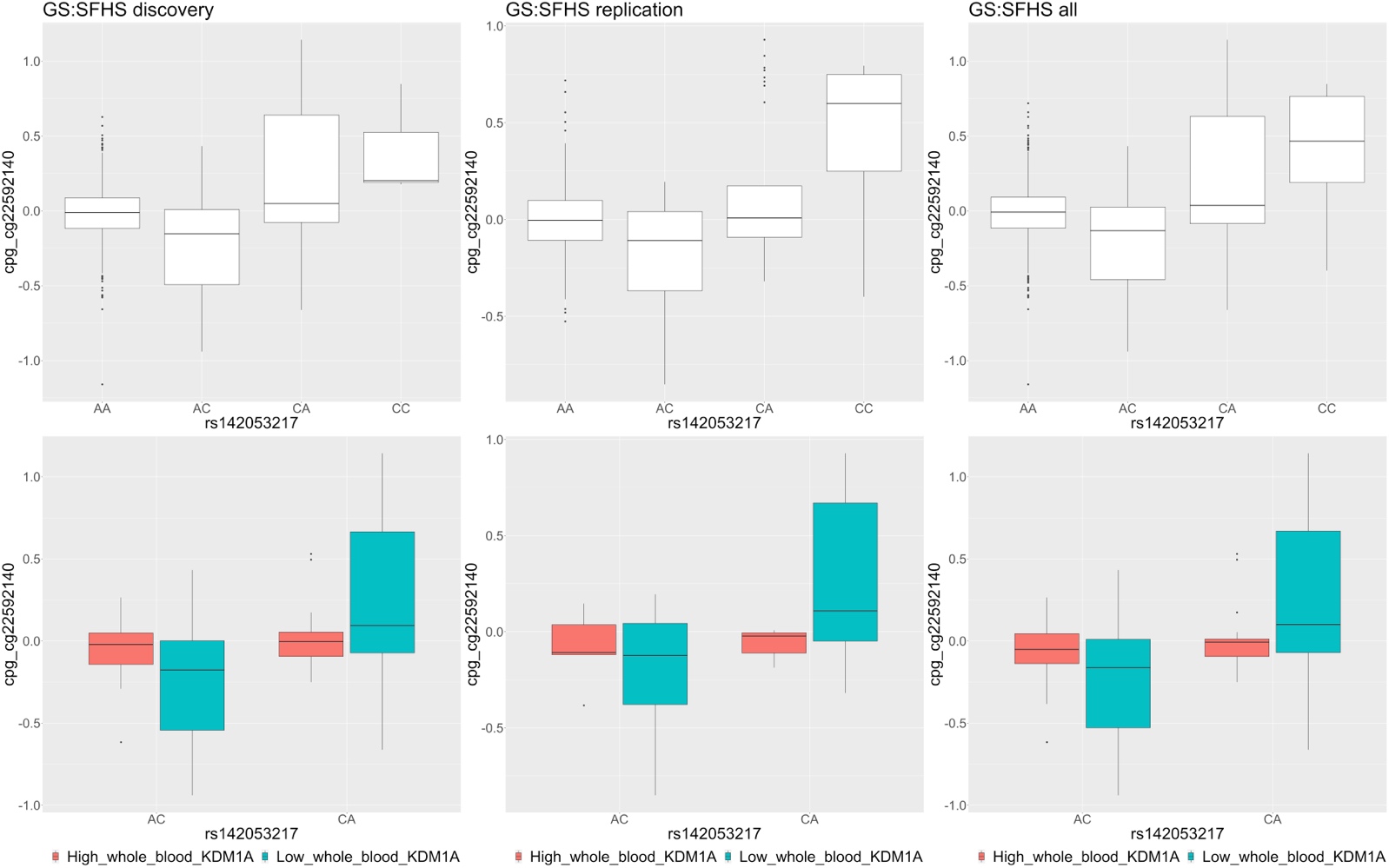

Figure s3. Correlations between cg21252175, ZNF609 (mRNA) and CLEC4C (mRNA) . a. The correlation between methylation level of cg21252175 and mRNA level of *ZNF609* in CD4+ peripheral T cells. b. the correlation between mRNA expression of *ZNF609* and the mRNA expression of *CLEC4C* in GTEX whole blood dataset. C. the correlation between mRNA expression of *ZNF609* and the mRNA expression of *CLEC4C* in single cell sequencing for PBMC cells of a healthy donor.

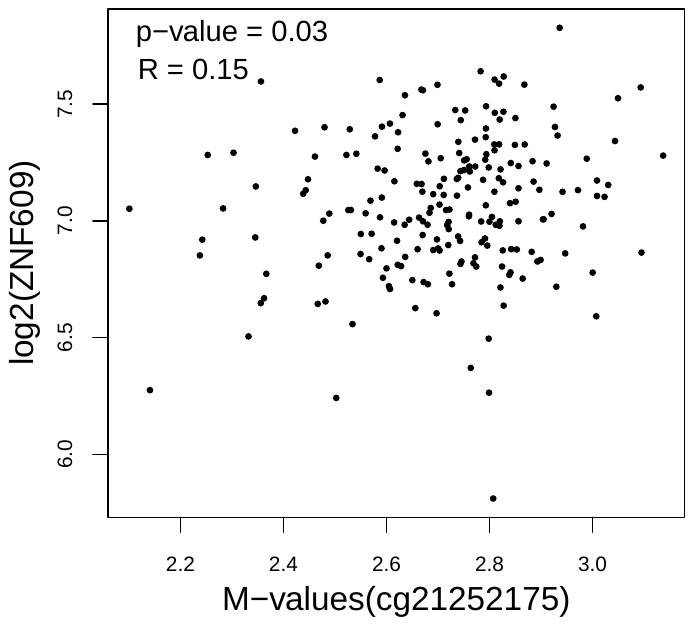
a

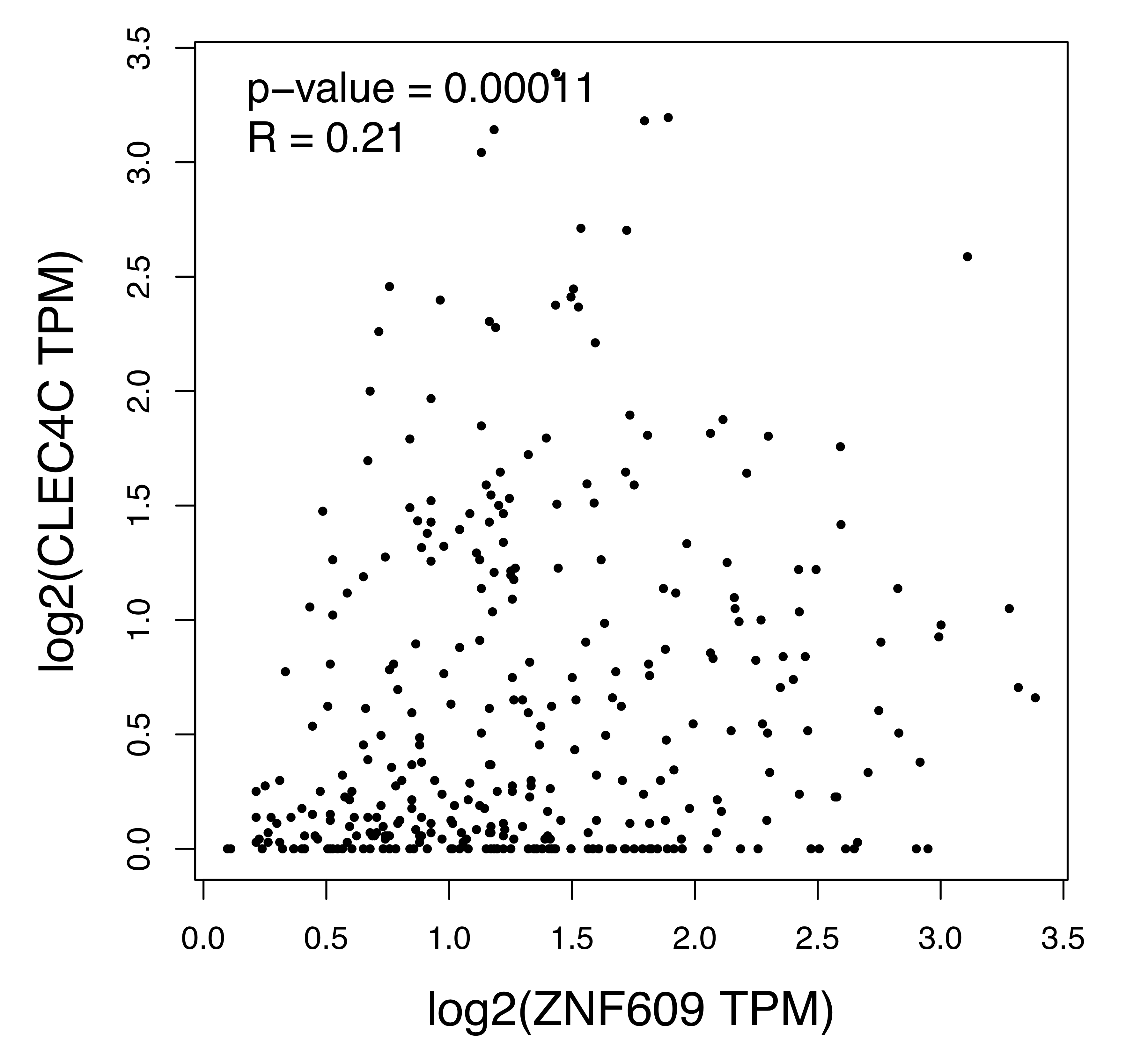

B

C.

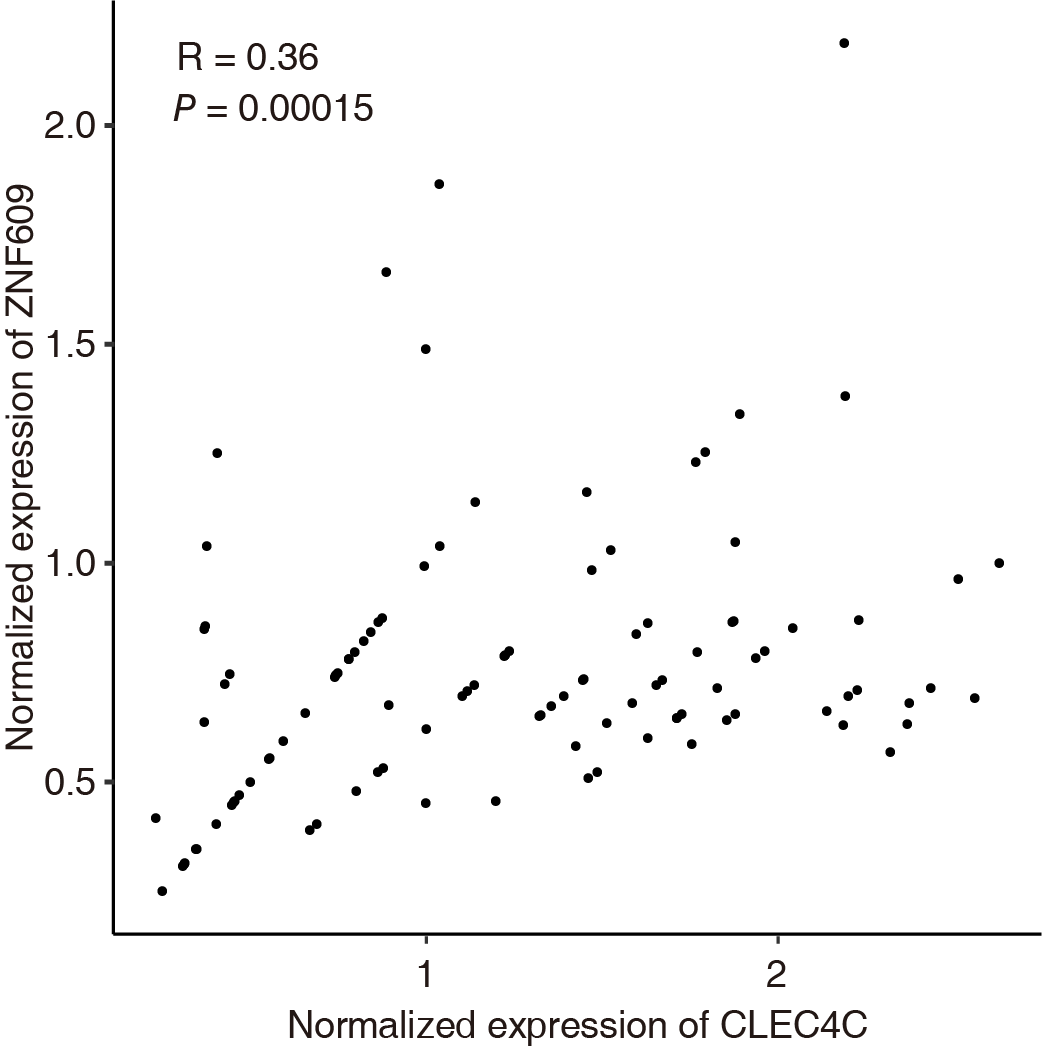

**Supplementary text 1**

**GS:SFHS: Quality control and pre-correction for DNA methylation data**

For each participant, whole blood DNA methylation was measured at 866,836 CpGs from using the Illumina Infinium MethylationEPIC array (<http://support.illumina.com>). Cell proportions were estimated for monocytes, granulocytes, B-lymphocytes, natural killer cells, CD4+ T-lymphocytes and CD8+ T-lymphocytes using the *estimateCellCounts* function in R package minfi[1]. Quality control was performed using the R packages shinyMethyl and meffil [2, 3]. Outlier probes and samples were removed based on signal intensity, control probe performance and consistency between registered and predicted sex. Samples with more than 0.5% of sites with a detection p-value > 0.01 were removed. Probes with more than 5% samples with a bead count smaller than 3 or overlapping with any common SNP (MAF≥0.01) in the European population in the 1000 Genome Project [4] were removed. The data were then normalized using the ssNoob method through R package minfi [5]. M values were obtained and adjusted as described below. Following previous protocols, we adjusted M values for potential technical variations including sentrix id, sentrix position, batch, clinic, appointment date, year and weekday of the blood extraction, and 20 principal components of the control probes through a linear mixed model [6]. To account for the genetic structure in GS:SFHS, we further fitted two random effects represented by genomic relationship matrices (G and K) to the technical-variation-adjusted methylation residuals [7]. The resulting adjusted M value residuals were used in statistical analyses.

**Supplementary text 2**

**Historic Scottish birth cohorts used to derive birth-related environmental variables**

Historic Scottish birth cohorts include the Aberdeen Children of the 1950s [8]; the Aberdeen Maternity and Neonatal Databank [9]; the Walker Birth Cohort [10]; and the Scottish Morbidity Records (SMR02 – the Maternity Inpatient and Day Case record, and SMR11 – the Neonatal Inpatient dataset) (Ndc.scot.nhs.uk). This allowed us to additionally collect/derive a number of birth-related environmental variables [11].

**Genetic modifier variables**

1) Predicted mRNA expression levels of DNA methylation or imprinting-specific machinery genes. Table s2 contains a list of 30 genes which we collated from the literature on the basis of their critical roles in DNA methylation in general, such as methylation writers and erasers, and in imprinting specifically. Since GS:SFHS does not have mRNA expression levels directly measured on participants, we used PrediXcan [12] to impute mRNA expression levels in whole blood for the selected machinery genes using models trained on measured whole blood expression levels in the GTEx consortium. PrediXcan provided prediction models for 18 of the 30 machinery genes. Apart from one gene (*ZFP57*) which did not have enough SNPs for the prediction, the rest of the genes (N=17, table s1) had their whole blood mRNA expression levels successfully imputed in GS:SFHS participants and were used in analyses.

2) Genetic scores for folate metabolism. Folate and one-carbon metabolism play important roles in DNA methylation dynamics and genomic imprinting[13]. Genetic variation can contribute to individual variation in this pathway. We first created three polygenic risk scores (PRSs) for dietary folate intake for GS:SFHS participants. Summary statistics of GWAS of dietary folate were collected from the OpenGWAS database [14]. Three dietary-folate-PRSs scores were created at P value cutoffs of 0.01, 0.2 and 0.5. Next, six PRSs for the folate pathway and one-carbon metabolism were created using GWAS statistics for levels of vitamin-B12 and vitamin-B6 [15], serum folate [16], and one-carbon metabolites (cystathionine, homocysteine, first principal component of multiple metabolites) [17]. Since we did not have access to the effect sizes of all SNPs in these GWASs, we only used the genome-wide significant SNPs to construct PRSs. In total, nine folate-related PRSs were constructed and used in analysis.

**Supplementary text 3**

**Accounting for multiple testing**

In the discovery stage, we applied multi-step permutation analyses to account for multiple testing (N_test=independent_trio_= 2372*101=239,572) performed in the discovery sample while keeping the computational burden at manageable level. In first step, a global-permutation analysis was applied to establish a global null distribution for the *POE_mQTL_ x Mod* interaction effect which allowed the estimation of a single significant threshold for the observed data at FDR≤0.05 level [6, 18, 19]. In detail, the interaction model was rerun for the entire dataset after shuffling individual identifiers for the methylation data and keeping identifiers for other variables unchanged. This allowed the construction of a null distribution while the correlation structure among mQTLs, CpGs and modifiers was maintained. Ten replicates were used following previous studies [6, 18, 19], and this led to the estimate of the FDR≤0.05 p-value threshold to be 2.93 x 10^-5^. A total of 1,088 independent mQTL-modifier-CpG trios (3,289 trios in total) reached this threshold. A secondary permutation test was then performed to estimate trio-specific permutation-based P values. In brief, for each tested trio the interaction model was rerun for 1x10^5^ times using permuted methylation data (N_permutation_=1 x 10^5^), which allowed the calculation of a trio-specific permutation P value after locating the observed P in the ranked P values from permuted data. The Bonferroni method was used to adjust for multiple testing (N_correction_=1,088 independent mQTL-modifier-CpG trios), 516 independent trios (1,878 trios in total) reached an adjusted P<0.05 and were deemed as significant in the discovery sample (Table s4).

In the replication stage, the same interaction model was applied to test for the *POE x Mod* interaction effects for the 516 candidate independent mQTL-modifier-CpG trios (identified in the discovery stage, 1878 trios in total) in the replication sample. FDR was applied to adjust for multiple testing correction (N_correction_=516) and 46 independent trios (69 trios in total) passed the adjusted significance threshold of P < 0.05. To ensure that a successful replication was only reported when the interaction effect was both statistically significant and consistent for both the POE (main effect) and the POE-related interaction effect across discovery and replication samples, we performed three additional filtering:

1) 39 independent trios (40 trios in total) were excluded because the main parent-of-origin effect was not statistically significant (Table s5) in the replication sample in a POE model with no interaction term (the model 1 in Methods). Model 1 was the POE-mQTL model which was used initially to discover POE-specific mQTL-CpG pairs in our previous paper[6]. Seven independent trios (29 trios in total) passed this first filtering (Table s4);

2) two independent trios (11 trios in total) were further excluded as the pattern of the interaction was not fully consistent between discovery and replication samples, five independent trios(18 trios involving four independent CpGs) passed this second filtering(Table s4);

3) one independent trio was additionally excluded as the CpG failed to reach Bonferroni-corrected significance in a CpG-level permutation test (N_correction=independent_cpg_=4, table s6). The test was to determine whether the lowest P value for the *POE_mQTL_ x Mod* interaction effect on a given CpG was more significant than expected by chance (N_permutation_=1 x 10^4^), which helped to rule out the possibility that significance in trios involving a given CpG was due to an average higher test statistic level under the data structure of that CpG. We considered trios that passed all of these filters as successfully replicated (four independent trios remained). Detailed information for trios that passed or did not pass the filters described at each stage is shown in table s4.

**Supplementary text 4**

**ORCARDES cohort: Quality control and pre-correction for DNA methylation data**

Quality control was performed using minfi and meffil R packages[1, 3]. Samples were removed if > 1% of probes had detection p-value > 0.01, if samples showed evidence of dye bias or samples were an outlier for either of the bisulphite conversion control probes, if there was discordance between recorded sex and predicted sex from DNA methylation data and due to the median methylated signal intensity that was more than 3 s.d. lower than expected. Probes were removed if they had bead count > 3 in at least 5% of samples or had detection p-value > 0.01 in more than 1% of samples. Additional probes were removed if they were flagged as cross-creative[20]. Prior to normalization, sample-probe pairs with detection p-value > 0.05 were removed. Normalization was carried out using preprocessNoob() in minfi R package. DNA methylation data was stored in two formats: 1) Beta values which range between 0 and 1 and represent the percentage of methylation at each CpG site, and 2) M values as logit transformed Beta values. Linear mixed modelling was used to correct the M values for the technical covariates including plate number as random effect and top ten principal components of control probe intensities, season of venepuncture, year of venepuncture and position on the plate as fixed effects. Residual M values were available for 794,627 CpG sites in 1052 samples. The estimateCellCount2 in minfi R package was used to estimate white blood cell proportions which were stored for later use in the analysis[1].
